# Towards the NMR solution Structure and the Dynamics of the C-terminal Region of APOL1 and its G1, G2 Variants with a Membrane Mimetic

**DOI:** 10.1101/2021.03.16.435683

**Authors:** Sethu M. Madhavan, Alexandar L. Hansen, Shufen Cao, John R. Sedor, Matthias Buck

## Abstract

Secreted apolipoprotein L1 (APOL1) is well known as an innate immune factor, protecting against African trypanosomiasis. The intracellular form has multiple functions, including regulating autophagy, intracellular vesicle trafficking, and ion channel activity. The APOL1 protein (G0) has two common variants (denoted G1 and G2) in the C-terminal region and are associated with a high risk of chronic kidney disease (CKD) and progression to end-stage kidney disease. Our previous studies using molecular modeling suggested that APOL1 G1 and G2 stabilize an autoinhibited state of the C-terminus, leading to impaired intracellular interactions with SNARE proteins. To characterize the structural consequence of kidney disease-associated APOL1 variants further, we assigned the C-terminal region proteins using ^1^H, ^13^C, ^15^N multidimensional nuclear magnetic resonance (NMR) spectra in solution in the presence of membrane mimetic dodecylphosphocholine micelles. We then derived models for the three-dimensional structure of APOL1-G0, and -G1 and -G2 variant C-terminal regions using the chemical shifts of the main chain nuclei followed by NMR relaxation measurements. The data suggest that changes in the three-dimensional structure of APOL1 C-terminal region induced by kidney disease-associated variants, not least the alteration of key sidechains and their interactions, could disrupt membrane association and the yet to be characterized protein-protein interactions including its binding partners, such as SNARE proteins. Such interactions could underlie the intracellular mechanisms that mediate the pathogenesis of CKD. In the future, one may try to reverse such structural and dynamics changes in the protein by designing agents that may bind and then mitigate APOL1 variant-associated CKD.

## INTRODUCTION

Common variants in the *APOL1* gene (termed G1: mutation S342G & I384M and G2: deletion of 2 residues, N388 & Y389, **Fig. 1A**) in a recessive genetic model are associated with the risk of chronic kidney disease (CKD) in individuals with African ancestry (1–4). In the healthy human kidney, the APOL1 protein is synthesized and expressed in podocytes, glomerular and extraglomerular endothelium, and circulates as a complex with HDL (5–7). The variant protein that is expressed in kidney cells itself is directly associated with the risk of CKD (8,9). Not all individuals with two copies of APOL1 variants develop kidney disease suggesting that an additional genetic, immune or environmental “second stress” stimuli is necessary to manifest clinically significant kidney disease (1). Studies aimed at understanding the molecular mechanisms by which APOL1 variants result in CKD pathogenesis and progression are evolving. These studies have shown that APOL1 variants dysregulate cellular homeostasis resulting in increased lysosomal permeability, loss of mitochondrial membrane integrity and cellular energy depletion, endoplasmic reticulum stress, and disordered vesicular trafficking (6,10–21). In principle, these studies suggest that a change in the protein structure-function relationship could be responsible for the increased CKD risk of the two APOL1 variants, G1 and G2 (22). The biological function of proteins is usually intricately linked to their three-dimensional structure and, in many cases, also their stability, if not specific dynamic fluctuations in the protein. As shown in other studies, gene variations resulting from single amino acid changes can have drastic effects on a protein’s structure, dynamics, and function (23–25).

**Figure 1:**
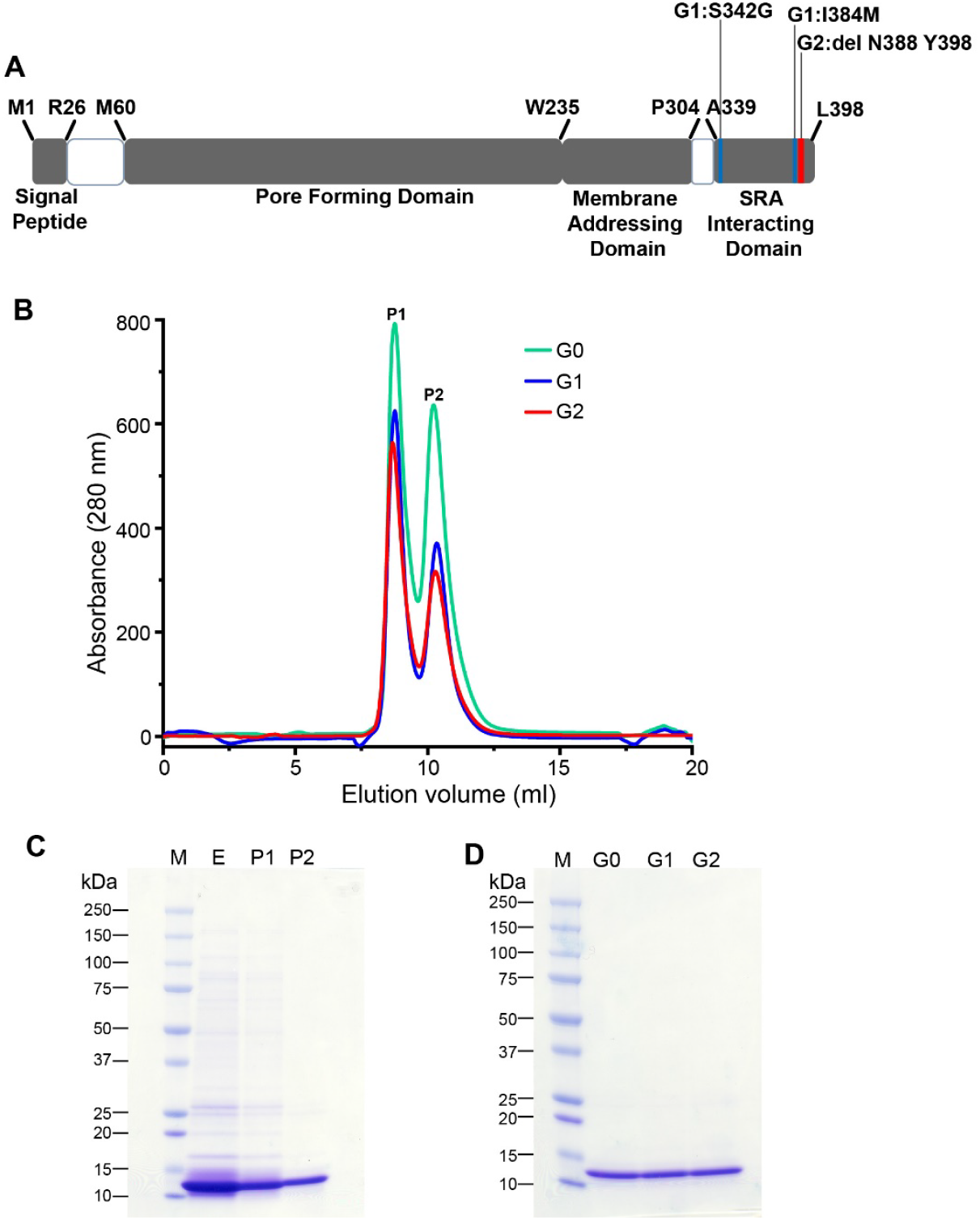
Domain organization of APOL1 and purification of APOL1 C-terminus (aa. 305-398) under non-denaturing conditions. (**A**) Schematic representation of APOL1 protein showing the different protein domains and location of G1 and G2 variants. The blue and red rectangle in the serum resistance associated (SRA) protein interacting domain represents location of the G1 and G2 variants, respectively. (**B**) Size exclusion chromatography profile of APOL1 -G0, -G1 and -G2 C-terminal domain purified under non-denaturing conditions in the presence of DPC micelles. Peak 1 is a large molecular weight state representing either protein oligomers or aggregates. Peak 2 is the monomeric fraction, which was used for CD and NMR studies. (**C**) Coomassie-stained SDS-PAGE gel of total eluate and peak 1 and peak 2 fractions from size exclusion chromatography of the wild type protein, G0. (**D**) Coomassie-stained SDS-PAGE gel of peak 2 showing purified APOL1 C-terminal domains of G0 and G1 & G2 domains (MW: protein molecular weight standard).

The protein structures of the reference APOL1 (G0) and of the G1 and G2 variants have not been experimentally studied. Previously, our group and others have used computational methods using template-based and *ab initio* modeling to characterize the structure of APOL1 protein domains (6,26–30). The domain organization of APOL1 is as follows (**Fig. 1A**): An N-terminal pore-forming domain (M60-W235) is predicted to adopt a helical bundle structure similar to bacterial colicins, and the membrane addressing domain (Q239-P304) is predicted to form a pH-sensitive helix (28,31). The C-terminal trypanosomal SRA interacting region of APOL1 is where the G1 and G2 variants are located (M329-L398) and this has been predicted to form an amphipathic helix (2,6,26,31). We previously modeled this C-terminal region of APOL1 using a protein structure predictive threading algorithm, followed by all-atom molecular dynamics (MD) simulations (6). These studies suggested that APOL1 C-terminus adopts an α-helical hairpin structure, and the presence of G1 or G2 variants appears to stabilize the secondary structure in the C-terminal region while impairing functional protein interactions. In this study, we present structural models for G0 and the G1, G2 variants, derived from experimental chemical shift data in solution NMR spectra. We found that the C-terminal region of APOL1 forms an amphipathic two helix bundle with relatively unstructured N-terminal residues in DPC micelles. The amino acid substitutions in the G1 variant likely result in the loss of a salt bridge (E355-K372) and altered interhelix hydrophobic contacts. In the G2 variant, the C-terminal region seems to adopt a more stable helical hairpin conformation by establishing additional, if not alternative interhelix hydrophobic contacts compared to G0. These changes are corroborated by effects on the internal dynamics of the mainchain, measured by ^15^N relaxation experiments, which is shifted locally in frequency, if not in amplitude. These observations add to our understanding of how the amino acid changes in the G1 and G2 variants result in an altered function of the proteins.

## METHODS

### Protein expression and purification

The cDNA for APOL1 aa 305-398 (NCBI reference sequence: NP_003652.2) was cloned into the NdeI and XhoI sites of the PET22b vector to generate pET22-*APOL1* (*305-398)-G0, -G1* and*-G2* constructs as previously described (6). This added eight extra amino acids (ELHHHHHH) to the C-terminus of APOL1 to assist protein purification. Protein expression and purification were modified, as detailed below, from a previously published protocol (6). To assist with the NMR assignments, four protein samples of the APOL1 G1 C-terminal region protein were also prepared with Val to Ala mutants: V346A, V349A, V353A, V346A+V349A+V350A, generated using Quikchange II Site-Directed Mutagenesis kit (Agilent Technologies, Santa Clara, CA). The plasmids were sequence confirmed and were transformed into *E. coli* BL21 DE3 cells for protein expression. Three to four colonies were picked from the LB plates and were inoculated into 10 ml of M9 medium and grown at 37°C overnight. Cells obtained from these overnight primary cultures were transferred to 1 liter M9 medium and grown at 37°C. For the preparation of NMR samples, ^15^NH_4_Cl (Sigma Aldrich, St. Louis, MO) and ^13^C-glucose (Cambridge Isotope Laboratories, Tewksbury, MA) were used as the sole nitrogen and carbon source in the M9 medium. Protein expression was induced with 0.75 mM isopropyl-β-D-thiogalactopyranoside at OD_600_ = 0.6 followed by incubation at 25°C for 20 hrs. Cells were harvested by centrifugation and were resuspended in 20 mls of lysis buffer (50 mM Tris, 20 mM imidazole, 200 mM NaCl, 0.5% ZA-14 detergent, 20 mM glutamic acid, 20 mM arginine, pH 7.4) supplemented with protease inhibitors (100 μM leupeptin, 10 μM bestatin, 20 μM pepstatin A, 200 nM aprotinin and 1 mM phenylmethylsulfonyl fluoride). The bacterial cells were lysed by sonication on ice (Branson digital sonifier 450) with a power level of 80% and a pulse length of 10 seconds on-off, using three 10-pulse-trains with a two-minute interval on packed ice. The lysate was then centrifuged at 16000x*g* for one hour at 4°C. The supernatant containing solubilized protein was filtered using a 0.22 μM nylon membrane filter and incubated with 2 ml of nickel-nitrilotriacetic acid (Ni-NTA) resin (Qiagen, Germantown, MD) at 4°C, which had been previously equilibrated with lysis buffer for one hour. Initial detergent screening for NMR was done using dodecylphosphocholine (DPC, Avanti Polar Lipids, Alabaster, AL), sodium dodecyl sulfate (SDS, Sigma-Aldrich, St. Louis, MO), n-dodecyl β-D-maltoside (DDM, Sigma-Aldrich, St. Louis, MO), n-octylglucoside (OG, Sigma-Aldrich, St. Louis, MO), 3-((3-cholamidopropyl) dimethylammonio)-1-propanesulfonate (CHAPS, Roche Life Sciences, Indianapolis, IN) micelles and also using 1,2-dihexanoyl-*sn*-glycero-3-phosphocholine (DHPC)/1,2-dimyristoyl-*sn*-glycero-3-phosphocholine (DMPC) bicelles (Avanti Polar Lipids, Alabaster, AL). Proteins bound to the Ni^2+^ column were washed with 40 column volumes (CV) of lysis buffer followed by buffer exchange to a wash buffer containing 50 mM Tris, 20 mM imidazole, 200 mM NaCl, 20 mM glutamic acid, 20 mM arginine solution also containing the membrane mimetic (containing one of 8 mM DPC, 0.4 % SDS, 5 mM DDM, 70 mM OG, 10 mM CHAPS or DHPC:DMPC bicelles with lipid: detergent molar ratio (*q*) at 0.5). Bound protein was eluted from the column in the 5 ml wash buffer with 500 mM imidazole. Eluted protein was concentrated with Amicon^®^ 3 kDa MWCO concentrator (Millipore, Burlington, MA) and injected into a 10/300 Superdex 75 size-exclusion chromatography (SEC) column (GE Healthcare Bio-Sciences, Pittsburgh, PA) equilibrated with SEC/NMR buffer (50 mM sodium phosphate, 100 mM NaCl, 20 mM glutamic acid, 20 mM arginine, separately with each detergent micelles or lipid bicelles at pH 6.8) and fractions of interest were collected for downstream applications. 2D-^1^H ^15^N-HSQC experiments were performed as described below to select conditions that gave the most cross-peak signals and best spectral dispersion.

### Circular dichroism (CD) spectroscopy

CD spectroscopy was performed under buffer conditions close to those of the NMR experiments. APOL1 proteins were purified by size exclusion chromatography to 50 mM sodium phosphate, 50 mM sodium chloride, 20 mM arginine, 20 mM glutamic acid, 5 mM DPC followed by the addition of DPC to achieve a final concentration to buffer containing 150 mM DPC. However, we observed a low signal/noise ratio between 225-190 nm in this condition. The sample was then buffer exchanged using a Sephadex G-25 column (GE Healthcare Bio-Sciences, Pittsburgh, PA) to CD buffer (20 mM sodium phosphate, 50 mM sodium sulfate, 5 mM DPC) containing 20 μM protein, which improved the signal to noise ratio below 225 nm. Far UV-CD spectra (190 nm to 260 nm) were recorded at 25°C in a quartz cuvette with 1 mm path length at varying pH (7.5, 6.8, and 5.0) on an Aviv CD spectropolarimeter. The pH of the sample was adjusted by the addition of o-phosphoric acid. Spectra were obtained with 1 nm step from 260 nm to 190 nm with a measurement time of 10 seconds at each wavelength. Background spectra were recorded for the CD buffer only and were subtracted from the corresponding spectra with the APOL1 protein. The secondary structure composition was estimated with Dichroweb using the CONTIN program (32–34).

### NMR spectroscopy: data acquisition, processing, and analysis

#### Structure determination experiments

^15^N and ^15^N/^13^C labeled APOL1 (aa 305-398) was concentrated to 0.6 mM in NMR buffer containing 50 mM sodium phosphate, 100 mM sodium chloride, 20 mM glutamic acid, 20 mM arginine at pH 6.8 with 150 mM DPC, 10 μM DSS (4,4-dimethyl-4-silapentane-1-sulfonic acid) and D_2_O at 10% v/v. All NMR experiments were performed at 313K on Bruker Avance 800, 900 MHz spectrometer equipped with TXI cryoprobes and 800 MHz Bruker Avance III HD spectrometer equipped with a 4-channel QCI cryoprobe. Pulse programs were obtained from the Topspin v3.5 pl7 pulse program library (Bruker, Billerica, MA). Chemical shift assignments were carried out using 2D-^1^H-^15^N-HSQC, 3D-HNCA, 3D-HNCO, 3D-HN(CO)CA, 3D-HNCACB, and 3D-CBCA(CO)NH experiments. Additional mutant APOL1-G1 proteins (V346A, V349A, V353A, V346A+V349A+V350A) were expressed, purified, and analyzed to facilitate assignments. Spectra were processed with TopSpin version 3.5 pl7 software (Bruker) and analyzed using NMRFAM-Sparky version 1.412 (35) and CARA (36). Backbone chemical shifts were used to indicate the secondary structure of the protein from TALOS-N (37), and models of the three-dimensional protein structure informed by the chemical shifts were obtained using CS-ROSETTA v3 (38). Figures of the structures were generated using Pymol version 2.0.4 (39). The assignments are deposited in BMRB under accession numbers 50685, 50675, and 50605 for G0, G1, and G2 C-terminal domains, respectively.

#### NMR relaxation experiments

^15^N relaxation (R_1_, R_1ρ_, [^1^H]-^15^N hetNOE) experiments were performed using ^15^N labeled C-terminus of APOL1 protein at a concentration of 0.6 mM in NMR buffer in 150 mM DPC, 10 % D_2_O and 10 μM DSS. All NMR relaxation experiments were acquired on an 800 MHz Bruker Avance III HD spectrometer equipped with a 4-channel QCI cryoprobe at pH 6.8 and 313 K. R_1_, R_1ρ_ relaxation, and hetNOE spectra were recorded with spectral widths of 12,820 Hz sampled over 2,048 complex points in the ^1^H dimension and 1,946 Hz over 128 complex points in the ^15^N dimension with 16 scans. Relaxation delays were 40, 400, 760 (x2), and 1160 ms (x2) for R_1_ and 2, 32, 62 (x2) and 92 ms (x2) for R_1ρ_. Duplicates of two relaxation times were done for all experiments for error estimations. Relaxation data were processed using NMRPipe/nmrDraw software (40) in NMRbox and graphed using the OriginPro version 2020. NMR spectra were referenced to the internal DSS signal. R_1_ and R_1ρ_ values were calculated by fitting peak intensities using the single exponential decay function *I(t) = I*_*0*_*exp(-R*_*1ρ*_*t*). R_2_ was calculated using the relation *R*_*2*_ = (*R*_*1ρ*_ – *R*_*1*_ *cos*^*2*^(*θ*))/*sin*^*2*^*(θ)*, with *θ* representing the effective field angle for each resonance and is determined from *θ = tan*^−1^*(ω/Ω)*, *ω* and *Ω* represent the spinlock power and resonance offset respectively. R_2_ error was estimated as:

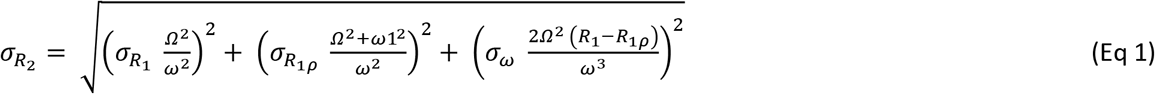

Heteronuclear [^1^H]-^15^N NOE values were measured without and with initial proton saturation with a saturation period of 6 s. To calculate the hetNOE, cross peak intensities were measured as peak heights, and errors were estimated from the standard deviation of the signal/noise ratios of individual peaks as *σ*_*NOE*_*/NOE* = ((*σ*_*Isat*_/*I*_*sat*_)^*2*^ *+* (*σ*_*Iref*_*/I*_*ref*_)^*2*^)^*1/2*^ where I_sat_ and I_ref_ represent the signal intensities of the saturated and reference spectra, respectively, and *σ*_*Isat and*_ *σ*_*Iref*_ represent the standard deviations of noise in the spectra.

## RESULTS

### Expression and purification of the APOL1 C-terminal domain

We have previously optimized a method for purifying the C-terminal domain of APOL1 from inclusion bodies under denaturing conditions using urea followed by on-column refolding (6). The kidney disease-associated APOL1 variants G1 and G2 are located in the C-terminal region of APOL1 (**Figure 1A**). To study the structure and dynamics of the APOL1 C-terminal domain here, we expressed residues R305-L398 of APOL1-G0 and the -G1 and -G2 variants in a bacterial system and purified the recombinant proteins under non-denaturing conditions. Our studies indicated that recombinant APOL1 C-terminal region protein has a high tendency to aggregate at the concentrations needed for solution NMR experiments unless membrane mimetic micelle or bicelle were added. After detergent screening, we found that the protein in dodecylphosphocholine (DPC) micelles gave the best ^1^H-^15^N HSQC NMR spectra with more uniform peak intensities and better dispersion compared to other detergents/bicelles (see next paragraph). Upon purification in the presence of DPC micelles, the purified APOL1 protein eluted in two peaks (**Figure 1B**). Peak 1 represents oligomeric or aggregated fraction of APOL1 C-terminal region protein along with co-purified nonspecific bacterial proteins, and peak 2 represents the nonaggregated fraction of the APOL1 protein (**Figure 1C and 1D**). Using this method, we were able to obtain 1.4 mg of unaggregated ^15^N/^13^C-labeled APOL1-G0 and -G1 proteins from one-liter M9 media culture. The yield of ^15^N/^13^C-labeled APOL1-G2 was slightly lower with one-liter culture in minimal media yielding 0.9 mg of unaggregated protein. The purified proteins were stable in the presence of DPC at 4°C for more than a month and at 40°C for more than two weeks, as shown by the NMR studies.

### NMR spectra of APOL1 C-terminal domain in DPC micelles

Initially, using the appearance of the APOL1 G1 protein in ^1^H-^15^N HSQC spectra, we screened detergent micelles and lipid bicelles including DPC, sodium dodecyl sulfate (SDS), n-dodecyl β-D-maltoside (DDM), n-octylglucoside (OG), 3-((3-cholamidopropyl) dimethylammonio)-1-propanesulfonate (CHAPS) and 1,2-dihexanoyl-*sn*-glycero-3-phosphocholine (DHPC) and 1,2-dimyristoyl-*sn*-glycero-3-phosphocholine (DMPC) bicelles. Protein samples in DDM and SDS gave reasonable quality NMR spectra (**Supplementary Figure 1**), but the best quality spectra were obtained using DPC micelles (**Figure 2**), showed 80 out of the 90 expected peaks in G0 and G1 and 84 out of the 88 expected peaks for the G2 C-terminal domain. To assist with assignments, particularly of G1, we generated multiple protein constructs with mutations of certain valine residues to alanine (V346A, V349A, V353A, V346A+V349A+V350A).

**Figure 2:**
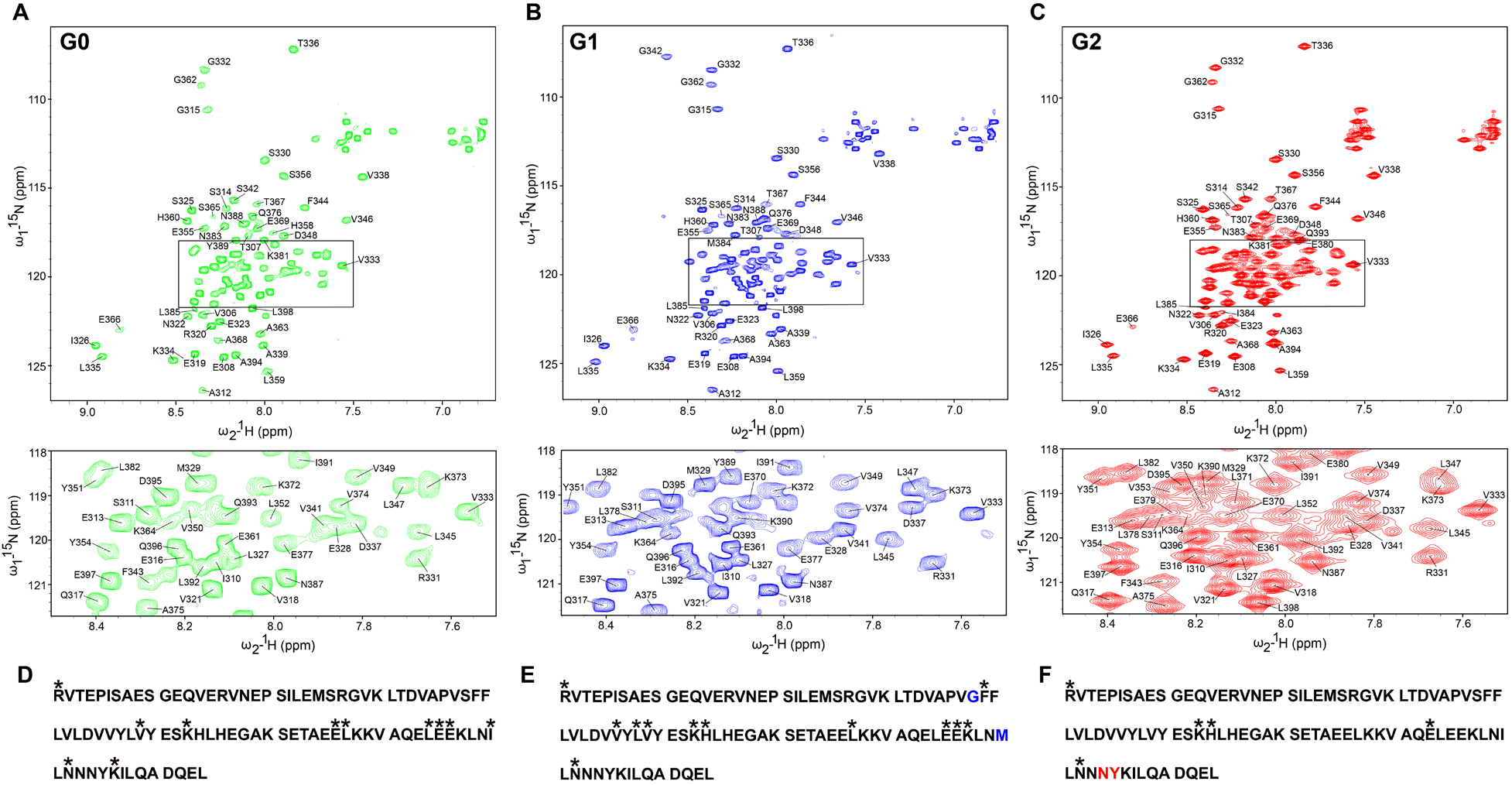
^1^H-^15^N HSQC spectra of APOL1 G0, G1, and G2 C-terminal domain in DPC micelles. (**A-C**) APOL1-G0, -G1 and -G2 C-terminal domain (aa R305-L398) at 800 MHz. Panels below, represent expanded regions from the corresponding HSQC spectrum (indicated there by a box) for better clarity. Residues, which occur in APOL1-G2 after the N388 and Y389 deletion, are numbered based on the G0 sequence. (**D-F**) Amino acid residues R305 to L 398 of APOL-G0, -G1 and ‒G2 are shown. Asterisk represents residues with missing H-NH assignments. Amino acid substitutions in G1 variant is shown in blue and deleted residues in G2 is represented in red.

### Secondary structure of APOL1 C-terminal domain in the presence of DPC micelles

Our prior studies evaluating the secondary structure composition of APOL1 C-terminal domain using far-UV circular dichroism (CD) spectroscopy in the absence of detergent micelles showed a predominant helical structure and no substantial difference in the secondary structure composition between reference and variant APOL1s (6). Also, the C-terminal region of APOL1 interacts with membranes and is predicted to undergo structural changes on lowering pH, mimicking the environment in late endosome and lysosome (27–29). We analyzed the changes in the secondary structure composition of the APOL1 C-terminal region in response to a low pH environment using CD spectroscopy in the presence of DPC micelles. This showed a similar α-helical content between -G0 and -G1 protein variants at pH 7.5 and 6.8, with an increasing α-helical content at pH 5.0, but with G1 significantly higher than G0. The C-terminal region protein APOL1-G2 exhibited a higher α-helical content compared to G0 and G1 over the entire pH range tested (**Figure 3A**). Additionally, ^1^H-^15^N NMR spectra of G0 C-terminus obtained at a lower pH range (pH 6.8-5.0) showed chemical shift changes consistent with pH-induced local conformational changes in the structure of the APOL1-C-terminal region proteins (**Supplementary Figure 2**). Importantly, even at a pH of 5.0 there is no indication that the protein’s tertiary structure is disrupted. The number of shifted resonances is roughly in accord with the 17 Asp+Glu residue sidechains, which may start to become protonated, following the earlier protonation of two histidines.

**Figure 3:**
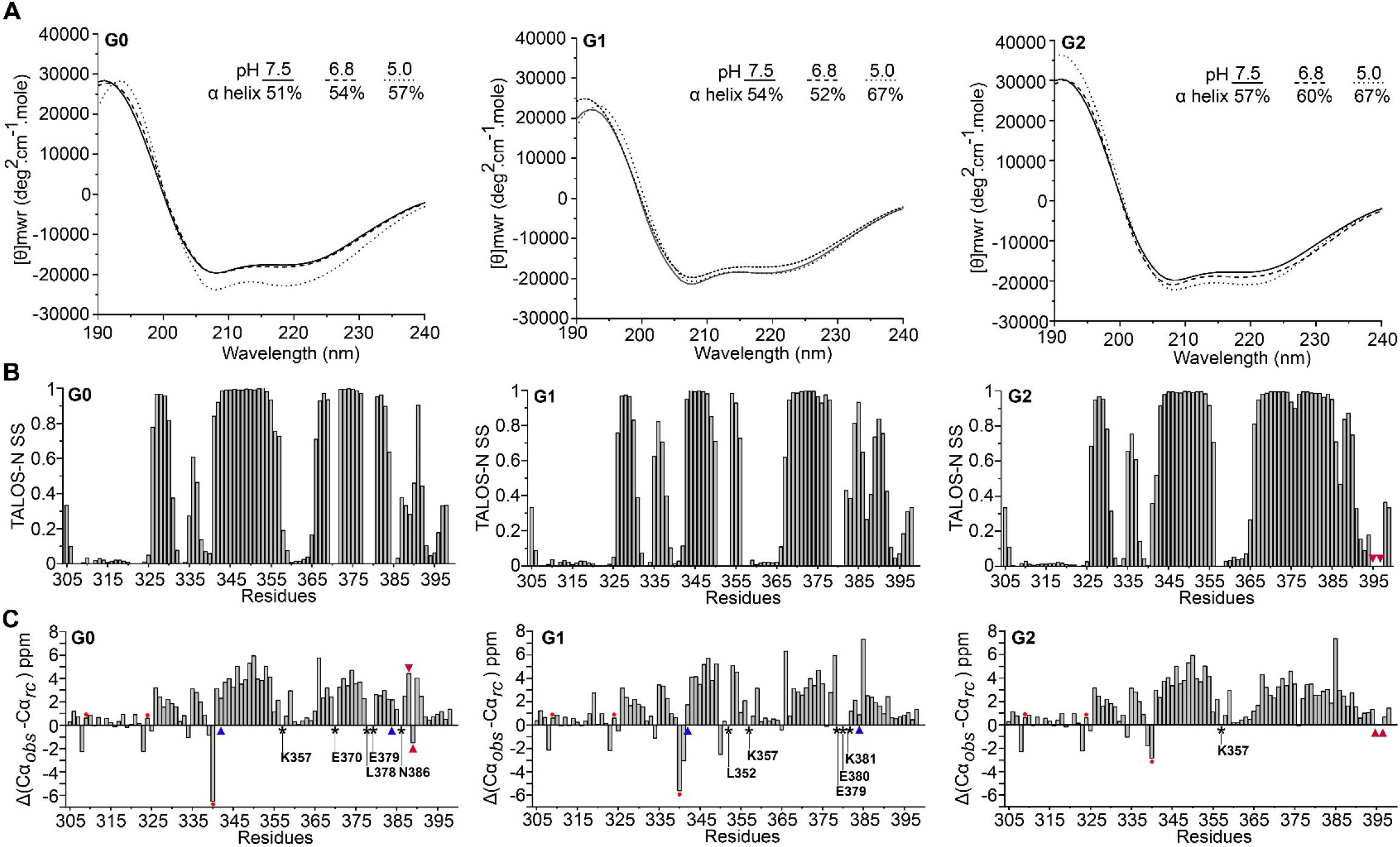
Secondary structure analysis of APOL1-G0, -G1, and -G2 C-terminal domains in DPC micelles. (**A**) CD spectroscopy of G0, G1 and G2 proteins in the presence of 5mM DPC micelles and at pH 7.5, 6.8, and 5.0. Calculated α-helical content at each pH is shown. (**B**) TALOS-N predicted secondary structure plotted as a function of sequence (**C**) Secondary structure determined using ^13^Cα chemical shift index for each residue. The residues with missing Cα assignments from the 3D-NMR experiments are highlighted by black asterisk above the corresponding labeled residues. Prolines in the sequence are marked by red dot. The location of the G1 and G2 variants are denoted by blue and red arrowheads, respectively.

Using 3D-NMR spectra, 89 out of the 94 Cα atoms were assigned for the G0 and G1 C-terminal domain. Chemical shifts of 91 for the 92 Cα atoms were assigned for the G2 variant. Using backbone chemical shifts obtained from NMR experiments, we analyzed the secondary structure of APOL1-G0, -G1, and -G2 with TALOS-N and also considered chemical shifts of Cα atoms by themselves (**Figure 3B-3C)**. This shows a predominant α-helical character of APOL1 -G0, -G1, and -G2 C-terminal protein region with the closely similar structural disorder at its N-terminal region (R305-S325). The two α-helical regions (I326-L359 and S365-L398) are separated by a short loop that is similar for all three proteins by Cα chemical shift and secondary structure profile. Overall, the C-terminal domain of APOL1 G2 showed more ordered α-helical conformation than the G0 and G1 variants but, except for the very C-terminal region (L385 to L398) this appears to be due to missing data in the G0 and G1, specifically around residue E370 (in G0), L352 (in G1), and residues L378-E380 (in both) (**Supplementary Figure 3C**).

### Models for the structure of the APOL1 C-terminal protein region in the presence of DPC micelles

We generated models for the three-dimensional structure of the C-terminal region of APOL1-G0, -G1 and -G2 using chemical shifts of the backbone atoms with CS-ROSETTA (**Figure 4 and Supplementary Figure 4**). Consistent with the secondary structure calculations, the N-terminal region (R305-P340) of the APOL1-G0, -G1, and -G2 C-terminal domain is mostly disordered. However, in many structures, a helical turn is seen around S330 and in some also around V338. This is consistent with secondary structure predictions and our previous modeling using sequence-structure threading (see discussion below). In the majority of the top 10 structures obtained for each protein, the N-terminal segment is attached to different regions of the protein (also see discussion). The remainder of the residues of the C-terminal region form two major α-helices (αH1 and αH2) with an intervening short loop, which results in a hairpin configuration of the two helices. The helices αH1 and αH2 extend over residues P340-E361 and S365-N388 in G0, P340-E361, and A368-I391 in G1, and V341-E361 and E366-L392 in G2 – thus, noting that the second helix is a turn longer in G1 and G2 (**Figure 4**). Amino acid variations induced by G1 and G2 altered the Cα-Cα and side chain inter-helix contacts formed between the αH1 and αH2. Although their sidechains point outwards, surprisingly, the S342G and I384M amino acid substitution in the G1 variant results in the loss of multiple interhelix (αH1-αH2) side chain hydrophobic contacts established by residues adjacent to S342 and I384M (P340-L385, V341-L382, V341-L385, F344-L378, F344-K381, F344-L382, F344-L385, and L345-L382) in the G1 C-terminus compared to G0 (**Figure 5**). Many of the low energy structures of the G1-C terminus showed a considerable crossing angle between the two helices, although some are also almost parallel but the helices are slightly rotated or shifted in G1 relative to G0, but one structure of G1 comes to within 4.4 Å of the lowest energy structure of G0. Noticeably, the salt bridge (E355-K372) that stabilized the helical hairpin in G0 and G2 is lost in the majority of low energy structures of G1, due to the helix rotation and/or sliding (**Figure 5**). By contrast, the deletion of residues N388 and Y389 in the G2 variant likely results in a stabilization of the helical hairpin by the formation of additional side chain contacts by multiple residues proximal and distal to the deletion compared to G0 (**Figure 6**). The deletion of these two residues located in the C-terminal region of the protein has made this second helix longer, and hence it is able to make additional interactions (**Figure 6**). This leads to sliding of helix-2 (αH2) (down a little) relative to helix-1 (αH1) and is accompanied by an altered helix-helix packing mediated by a flip of F344 (around it’s Cα-Cβ bond) and different hydrophobic contacts in G2 (V341-I381, F344-L382, F344-L385, F344-N386, L347-L378, L347-L382, L347-L385) compared to G0 (P340-L382, V341-L382, F344-L385, L347-L378) (**Figure 7**). The comparison of energies derived from the ten lowest energy CS-ROSETTA structures show that the G2 C-terminal region has slightly better energy compared to G0, but the G1 structures have on average a significantly worse energies (**Figure 8**).

**Figure 4:**
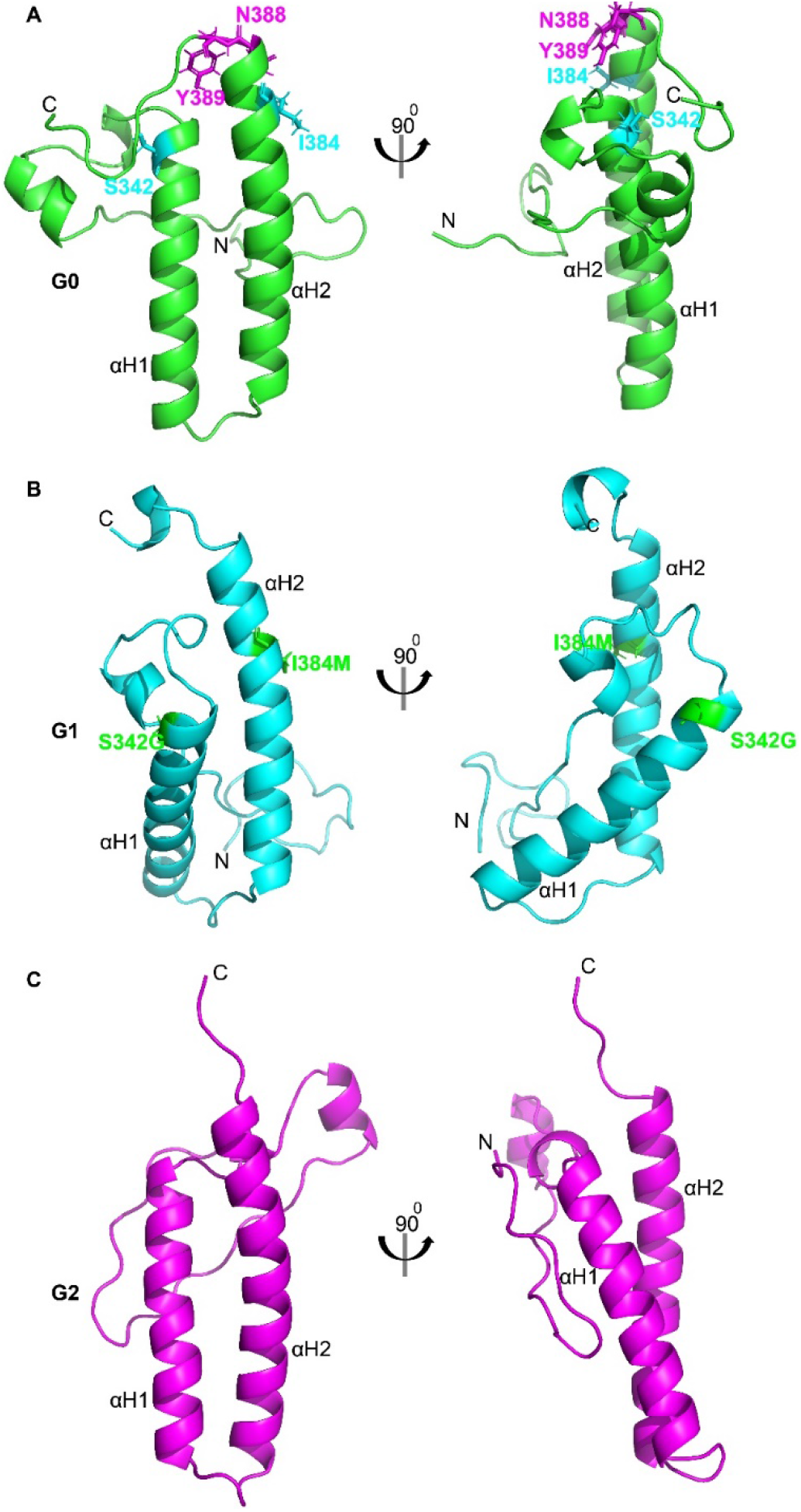
Models of structure for APOL1 C-terminal regions G0, G1, and G2 in presence of DPC micelles. (**A-C**) Lowest energy structure of the domain (aa: R305-L398) out of the 40,000 structures generated by CS-ROSETTA. The location of G1 (S342G & I384M) and G2 (two residue deletion of N388 & Y389) variants are highlighted (in case of G2 this is done in the G0 structure). αH1 and αH2 represent the two major α-helices. (N: N-terminus and C: C-terminus).

**Figure 5:**
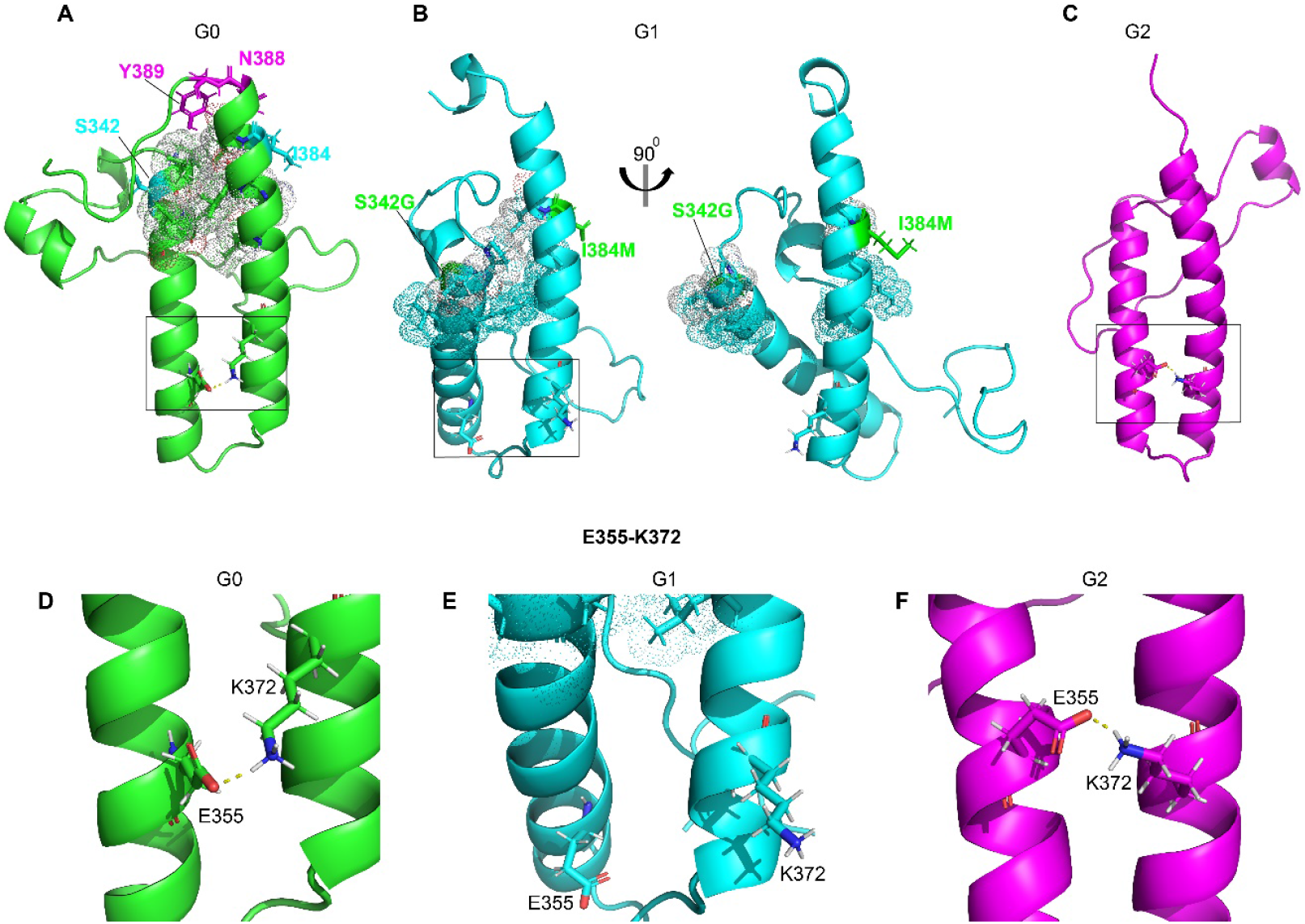
Loss of interhelix (αH1-αH2) hydrophobic contacts and salt bridge in the G1 C-terminal region. (**A**) G0 C-terminal domain structure showing the interhelix (αH1-αH2) hydrophobic contacts (represented in sticks and dot format) adjacent to the G1 variants S342G and I384M (represented in cyan). The interhelix salt bridge established between residues E355 and K372 is shown in yellow dotted line. The location of G2 variants is represented in magenta. (**B**) G1 C-terminal region showing loss of hydrophobic contacts adjacent to S342G and I384 M residues and of the interhelix salt bridge. Notice that the helices are rotated such that the sidechains point to opposite sides of the structure. (**C**) The C-terminal region of G2 showing the location of E355-K372 contact. (**D-F**) Salt bridge (denoted in dashed yellow lines in G0 and G2) formed between E355 and K372 stabilizing the helical hairpin is lost in the G1 C-terminal region compared to G0 and G2.

**Figure 6:**
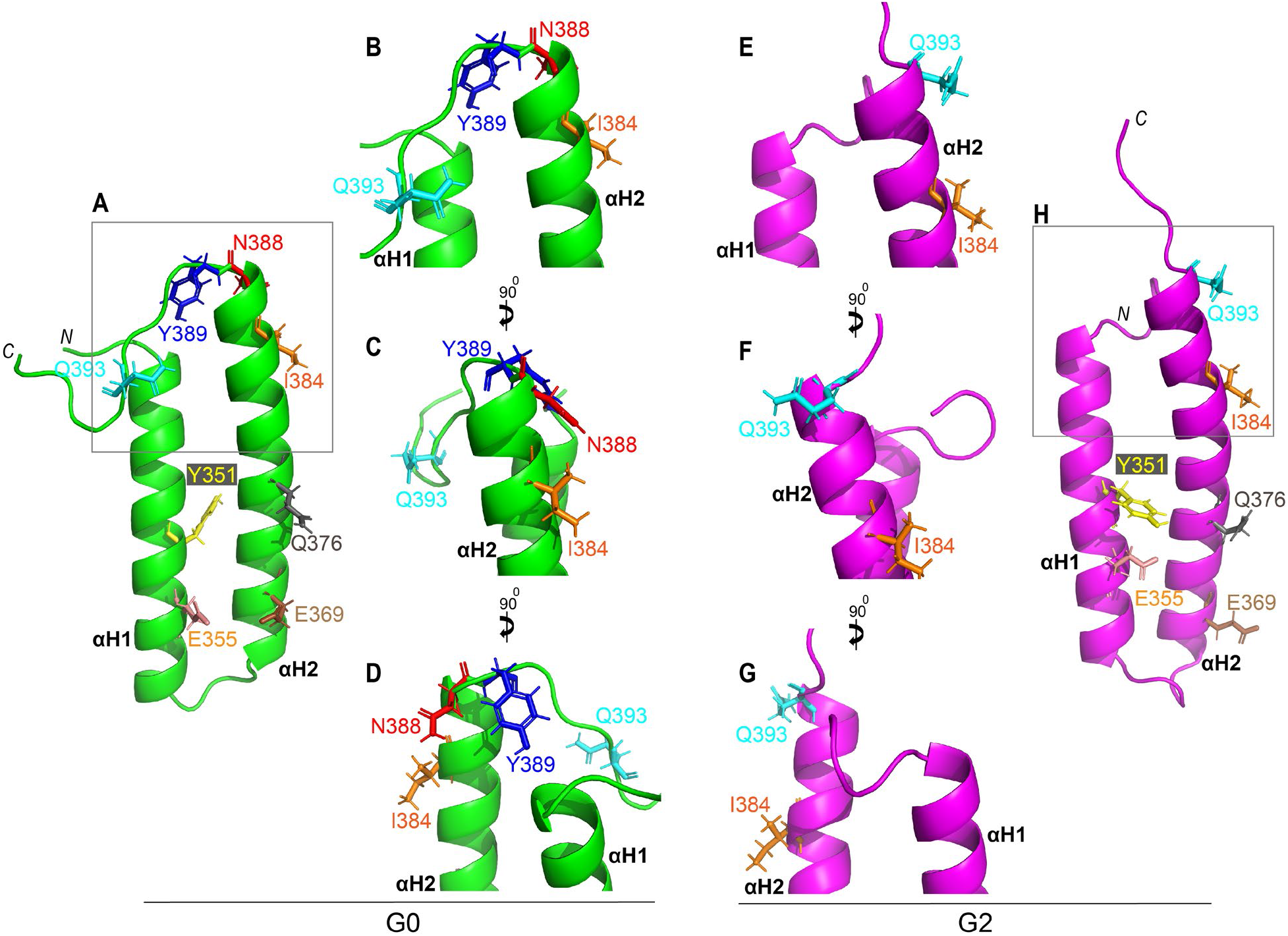
Extension of C-terminal helix in G2 variant. (**A-H**) NMR structure model of C-terminal αH1 and αH2 helices (residues D337-L398) from G0 (green) (**A-D**) and G2 (magenta) (**E-H**) obtained from NMR chemical shifts. Residues Y351, E355, E369, Q376 and I384 are highlighted in stick representation as reference which shows similar orientation in both G0 and G2. Residue Q393 (cyan) distal to the two deleted residues in G2 (N388 and Y389 represented in red and blue respectively) forms part of the unstructured region in APOL-G0 but remains in the helical region of G2 C-terminus. *N* and *C* represents N and C-terminus of the protein.

**Figure 7:**
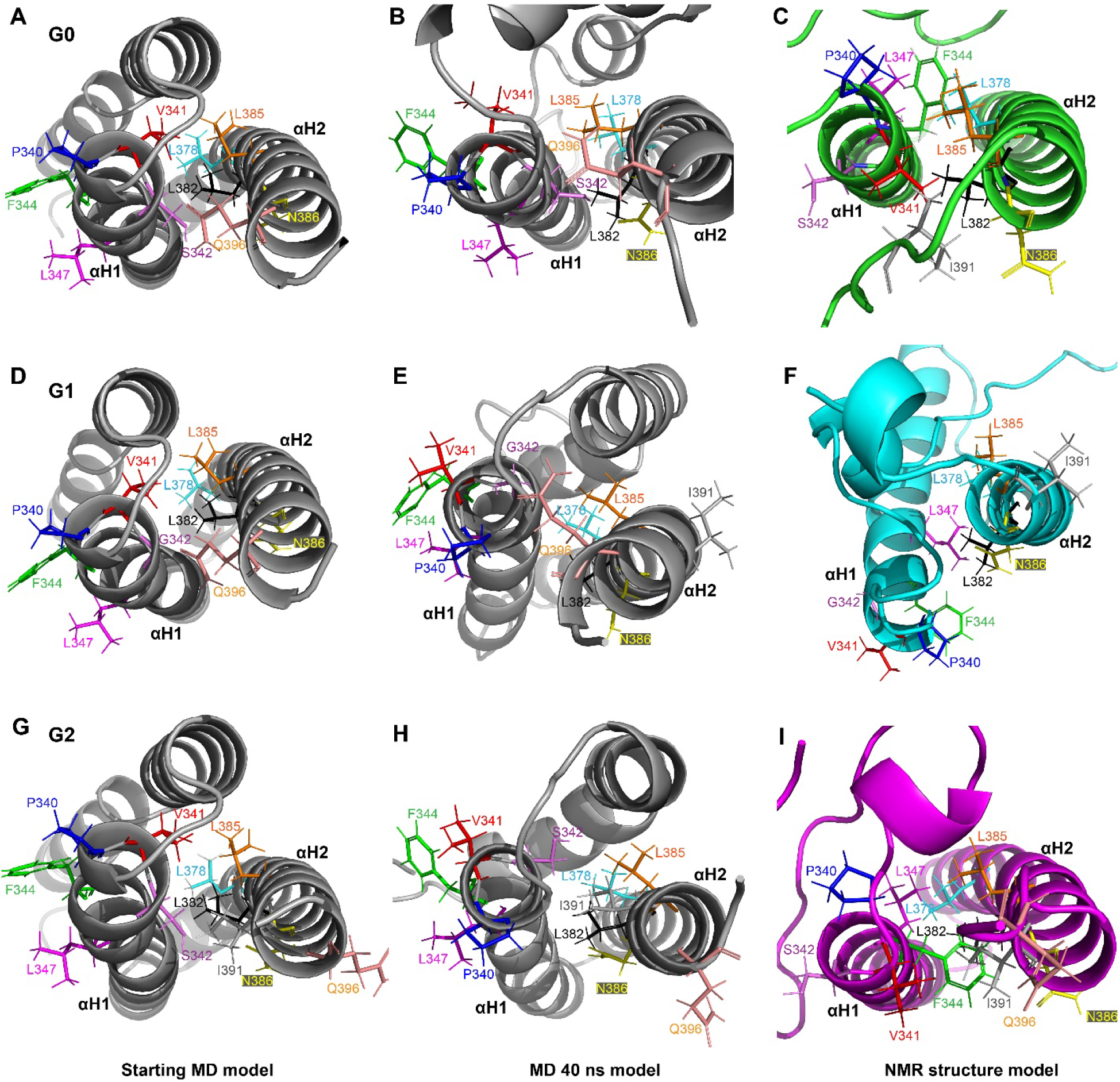
Comparison of helix packing of APOL1-G0, -G1 and -G2 C-terminus. The C-terminal helices of APOL1-G0 (**A-C**), G1 (**D-F**) and G2 (**G-I**) before and after 40 ns MD simulations compared to the lowest energy structure model obtained from NMR derived chemical shifts in CS-ROSETTA is shown. Selected residues in αH1 and αH2 are shown in stick representation to highlight the relative orientation of the helices. Amino acid substitution in G1 (I384M) and deletion of N388 and Y389 residues in the G2 variant of APOL1 results in extension of the αH2 compared to G0. This is accompanied by a slight rotation and sliding of the αH2 helix, relative to αH1, resulting in altered interhelix hydrophobic contacts and helix packing in the G1 and G2 C-terminus (**B and C**) compared to G0 (**A**). N and C represents N and C-termini of the protein.

**Figure 8:**
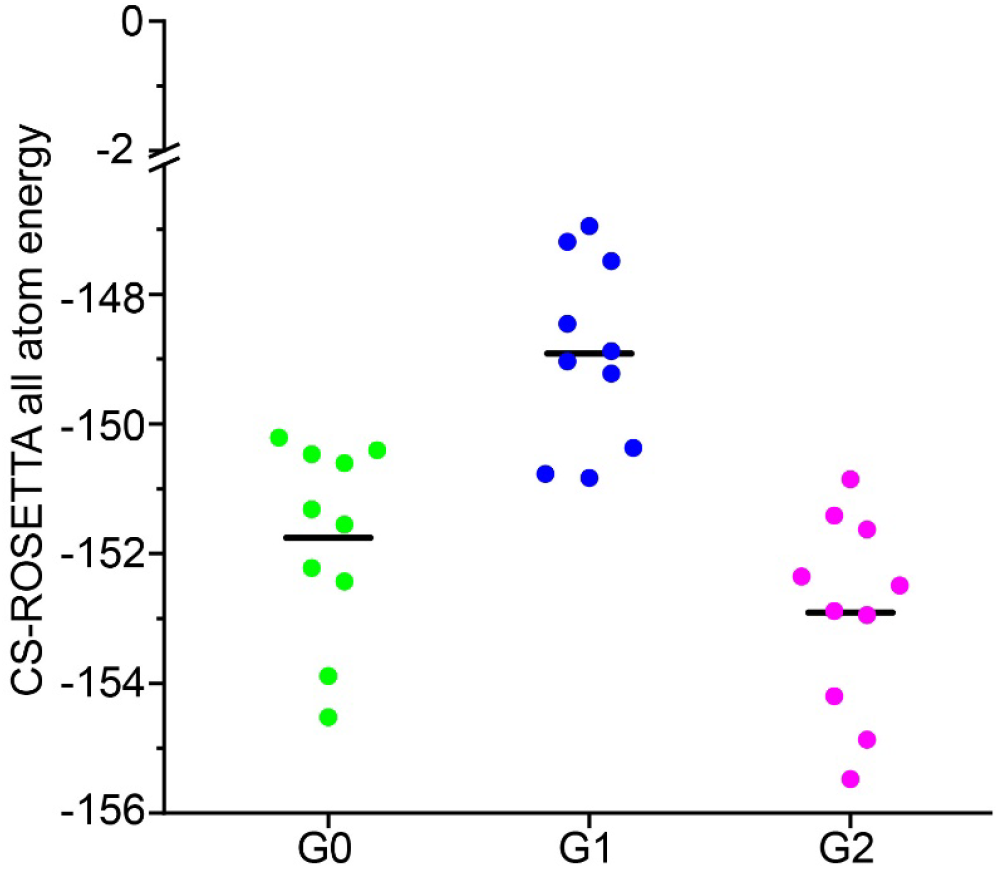
Comparison of CS-ROSETTA all-atom energy profiles of APOL1-G0, -G1, and -G2 C-terminal domain. All-atom CS-ROSETTA energy of ten lowest energy structures (out of the 40,000 structures) for APOL1-G0, -G1, and -G2 C-terminus. (Line denotes mean)

We compared the three-dimensional structure of APOL1-G0 obtained by solution NMR studies with the structures we previously predicted by a sequence-structure compatibility method (the “threading algorithm” iTASSER), and refined by all-atom MD simulations (6). The overall RMSD between the solution NMR structure of APOL1-G0 C-terminal domain (R305-L398) and pre-MD and post-MD (40 ns) *ab initio* predicted structures was 10.78 Å and 10.53 Å, respectively. This structural divergence was largely driven by the substantially disordered N-terminal residues (R305-P340) in the structures in presence of DPC micelles. However, a comparison of the region of residues A339-L398, which predominantly comprise α-helices αH1 and αH2 show lower RMSD values (pre and post-MD RMSD of 4.58 Å and 4.28 Å respectively) for G0, but this is still substantial, as the *ab initio* prediction generated structures with three helices (**Figure 9**).

**Figure 9:**
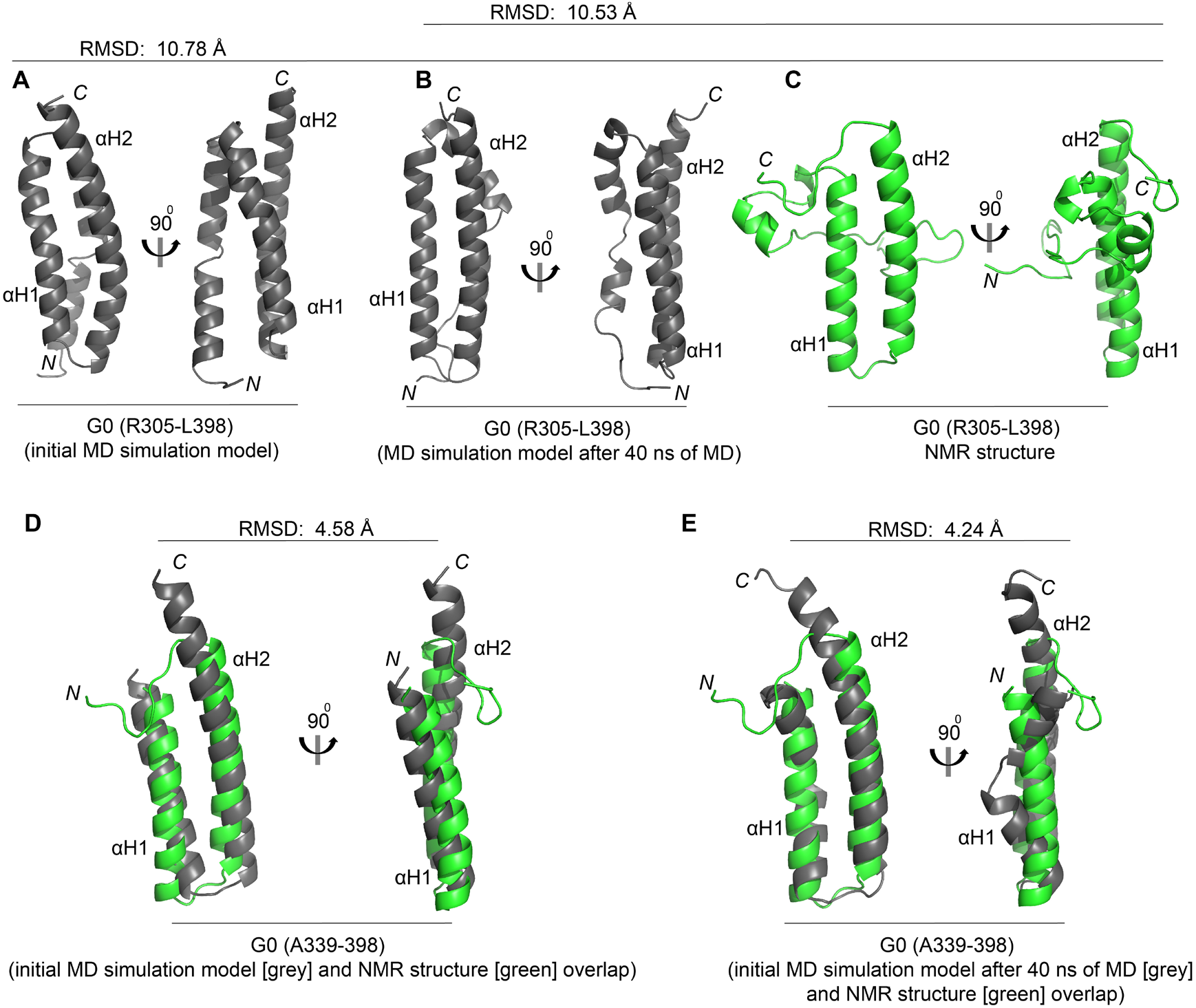
Comparison of the NMR derived model structures of APOL1-G0 C-terminal domain with the previously modeled structures. (**A-C**) Comparison of computationally modeled structure of APOL1-G0 C-terminus from I-TASSER server in grey (A) and the structure obtained after 40 ns of all-atom MD simulation in grey (B) with the NMR data derived structure in green (A305-L398) (C). (**D-E**) the NMR data derived structure (A339-L382) overlapped with initial computationally modeled structure (D) and after 40 ns of MD simulation (E). *N* and *C* represents N and C-terminus of the protein.

### Backbone dynamics of APOL1 C-terminal region in DPC micelles

The protein backbone dynamics that occur for main-chain amide bonds in the APOL1-G0, -G1, and -G2 C-terminal region proteins were probed using ^15^N-relaxation experiments, including R_1_ and R_2_ relaxation rates and [^1^H]-^15^N hetNOE experiments (**Figure 10**). The R_1_ and hetNOE measurements report on fast ps-ns motional amplitudes, whereas R_2_ also reports on μs-ms timescale motions. Except for a few isolated residues in G0 and G1, the N-terminal region appears to be highly flexible until about residue S325. It is interesting that the dynamics of the regions around residues S330 and V338 is similar in all three proteins, regions which have some helical turn character in the structures. In the case of G0, the R_2_ values are consistently slightly higher in the region L335-P340 compared to G1 and G2, suggesting that this part of the protein may be sampling several structures or contacts with αH1 and/or αH2 on the chemical exchange timescale. In fact, several residues in αH1 in G0 also shows slightly higher R_2_ values. The α-helical regions αH1 and αH2 in the APOL1-G0,-G1, and -G2 C-terminal domains showed low R_1_ and high R_2_ relaxation rates, and high hetNOE value suggesting rigid structures with relatively small amplitude conformational dynamics in solution (**Figure 10**). The inter-helix turn shows considerable local ps-ns flexibility in all three proteins and another segment, towards the end of the helix-2 (αH2) also shows similar large-amplitude ps-ns fluctuations. Finally, there is a systematic difference between residues K390-A394, with the extended helix-2 (αH2) in G2 undergoing some higher amplitude fluctuations on the ps-ns timescale. It is likely that N-H groups in all three proteins are affected by helix-coil transitions or by a collective fluctuation of this segment, but those transitions are faster and thus detected by the relaxation measurement in G2 (tighter structures leading to faster vibrations, shifted into the ps-ns timescale).

**Figure 10:**
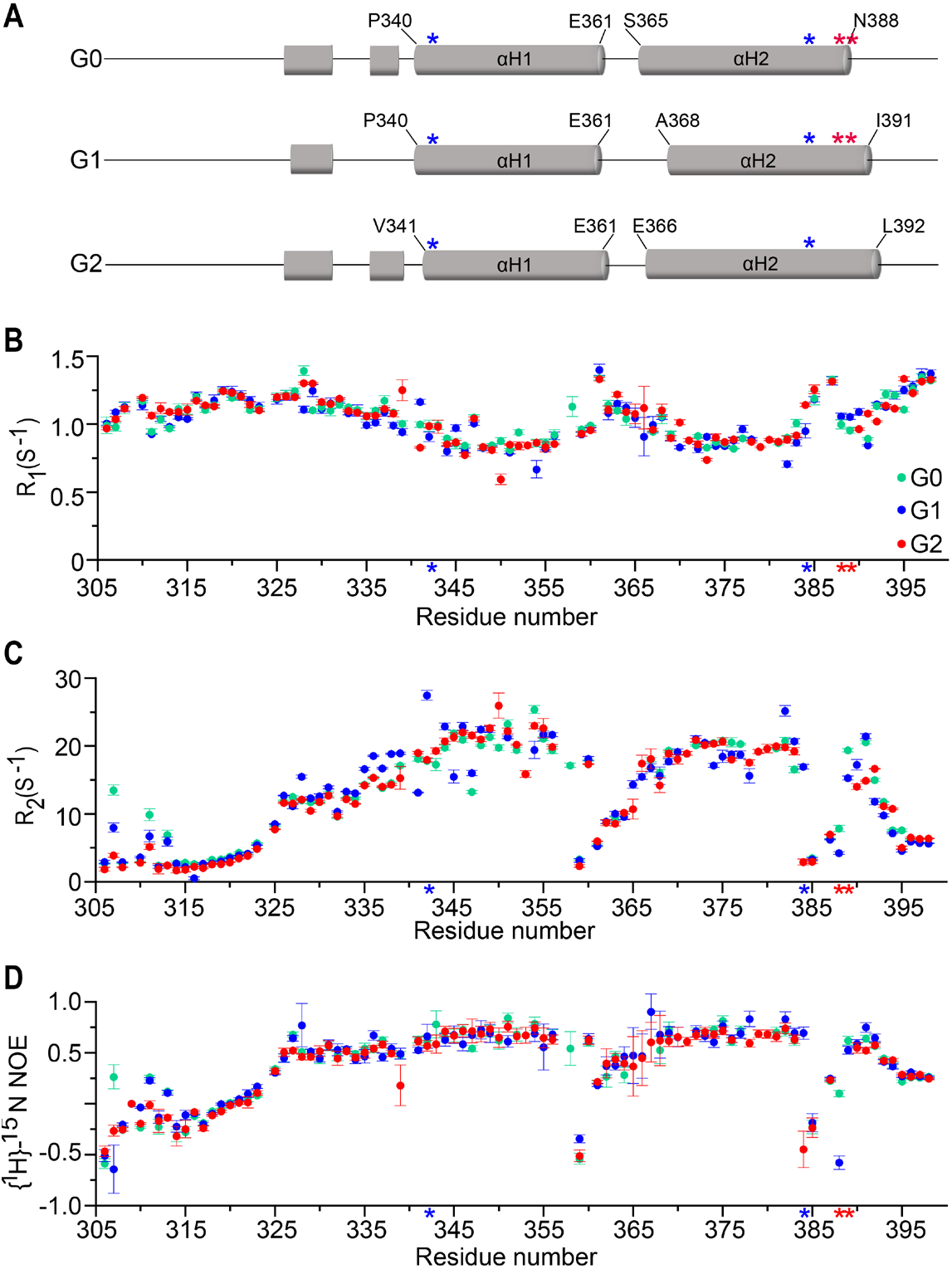
Backbone amide dynamics measurements by NMR relaxation of APOL1-G0, -G1 and -G2 C-terminal region in presence of DPC micelles. (**A**) Secondary structure representation of the C-terminus of APOL1-G0, -G1 and -G2 obtained from NMR chemical shifts. The location of G1 and G2 variant residues are marked by blue and red asterisk respectively. (**B-D**) Backbone dynamics demonstrated by ^15^N R_1_ (**B**), ^15^N R_2_ (**C**), and [^1^H]-^15^N het-NOE (**D**) values for APOL1-G0 (green), -G1 (blue) and -G2 (red) C-terminal domains recorded at 800 MHz are plotted against the residue number.

## DISCUSSION

Expanding on our previous study, which characterized the structure of the C-terminal domain of APOL1 using computational modeling and all-atom molecular dynamics (MD) simulations (6), we derived models for the three-dimensional structure of APOL1-G0, -G1, and -G2 C-terminal region (aa: R305-L398) using chemical shifts from solution NMR spectroscopy. The C-terminal domain of the wild type and variant APOL1s were purified under non-denaturing conditions in the presence of lipid/detergent micelles. After detergent and lipid bicelle screening, we chose DPC micelles as the detergent for our experiments, which has been used to solve structures of multiple other proteins using solution NMR. As predicted computationally in previous studies by our group and others (2,6,26,27,41), our solution NMR studies confirm that the C-terminal region of APOL1-G0, -G1 and -G2 formed an amphipathic α-helical hairpin with a largely disordered N-terminal region. The NMR derived models suggest that amino acid changes in the kidney disease-associated APOL1-G1, and -G2 variants induce moderate but overall considerable changes in the crossing angle if not the rotational orientation of the helices in the domain of APOL1 compared to the reference, the G0 wild type protein. The amino acid substitutions in G1 resulted in the loss of multiple interhelix (αH1 and αH2) side chain contacts adjacent to S342G and I384M, causing a change in the orientation of the C-terminal α-helical hairpin compared to G0. Consistent with the chemical shift data, relaxation measurements, and secondary structural content estimated from far UV circular dichroism, the two amino acid deletion towards the C-terminus of the G2 variant have an extended and likely stabilized αH2 compared to G0.

It should be noted that here we studied the protein in the presence of DPC detergent micelles. A noticeable difference is the reduced average linewidths in the center of the HSQC spectrum of the G1 protein, whereas G2 has more broadened peaks. This difference, although not manifest in the relaxation data due to the different time-scale involved, -suggests differences in solution behavior, possibly a chemical exchange in G2 (associated with the lengthened helix or a switch between helix-helix contacts) or a transient aggregation of the protein, if not different interactions with the detergent. Regarding interactions with DPC, this is the major difference between the current and our previous study (6), necessitated here by the need for higher protein concentrations for NMR experiments. The interaction with detergent, especially predicted for the N-terminal region of αH1 (V338-Y354) due to its hydrophobic nature could disrupt interactions with other helices, which was inferred from sequence-structure compatibility in our previous report. Although this helix was predicted to interact with both, what is called helix-1 (R305-A339) and helix-3 (S365-N388) in our previously published computational model, molecular dynamics refinement showed some distortion and partial unfolding. Here, in the presence of DPC only two short segments, near the N-terminus of the C-terminal APOL1 region (from the computational model) are modeled as helical turns, while the entire N-terminal region is making contact with either the front or more rarely the back-side of the helix-turn-helix structure. The relaxation data show that the N-terminal segment, prior to S325 is highly flexible, but this still leaves residues I326 to P340 to explore different regions of the highly folded part of the G0, G1 and G2 proteins. Remarkably, the relaxation and chemical shift data are very consistent in this region, with the exception of G0 in region L335-P340 as noted above and the overall impression of different linewidths particularly in the center of the HSCQ spectra of G1 and G2 (in opposite directions) compared with G0. In case of G1, residue S342 has been changed to G which may give this N-terminal region enhanced flexibility or ability to form interactions with the remainder of the protein, either of which could result in the lower linewidths. The alteration in the interhelix orientation will lead to changes in solvent-exposed residues in G1 and G2 variants compared to G0, which likely results in different orientational preferences of the domain with respect to a membrane surface. Use of the detergent, DPC only provides a very rough approximation and the presence of a more extended and planar membrane model system, such as a bicelle or nanodisc, is likely to enhance the differences between the reference and variant proteins. However, we also noted that the sidechains of the residue involved point outwards and are not directly involved in helix-helix packing interactions. In that sense, our previous model, where the G1 and G2 variants present a more stable autoinhibited structures in solution is not impacted by the findings of this report. Similar to our study, computational molecular modeling and MD simulation by Vanhamme *et al.* showed that the residues P340-L392 of APOL1-G0 formed an α-helix, which was then used to model the interactions with the trypanosomal SRA protein (27). The structure of the SRA protein has recently been determined as that of a helical bundle, suggesting an extended interaction surface with both αH1 and αH2 of the C-terminal region of APOL1 (42). Also, similar to our previous study, the models proposed by Tzur *et al.* and Sharma *et al.* predicted an α-helical hairpin structure for the C-terminus (S342-L398) of APOL1-G0, G1 and G2 (2,26). Sharma *et al.* also performed 2D-NMR studies on the C-terminus of APOL1-G0 and -G1 C-terminus (26). It should be mentioned that the overall ^1^H-^15^N-HSQC spectra previously published by Sharma et al., albeit for a shorter protein construct (A339-L398) in pure aqueous solution, are different from ours as many ^1^H-^15^N resonances are seen with up-field chemical shifts, suggesting a β-hairpin or β-sheet structure. In the absence of chemical shift assignments, it is even difficult to confirm whether or not APOL1 or another, possibly contaminant protein, was studied. For example, the ^1^H-^15^N HSQC NMR spectra in their study showed 83 and 93 peaks for the G0 and G1 protein constructs, respectively, compared to the expected 79 peaks. By contrast, we carried out resonance assignments and the chemical shifts of our spectra to derive a helix-hairpin structure overall consistent with previous computational predictions. Moreover, the mutations of the G1 and G2 variants showed multiple peak shifts in the C-terminal domain of APOL1, which were not limited to the variant residues (**Figure 2 and Supplementary Figure 3**), implicating non-local conformational changes.

The membrane addressing domain (Q239-P304), which is located N-terminal to the region we studied and the C-terminal region of APOL1 are predicted to undergo a large conformational change in the presence of a low pH environment (27,31). In an acidic extracellular environment that mimic the interstitial compartment of kidney, the APOL1-G1 and -G2 variants exacerbate mitochondrial structural abnormalities in HEK293 cells (43). We observed that the helical content of the C-terminus of wild type and variant APOL1s increased slightly in response to lowering pH with G2 C-terminal domain showing a higher helical content at the lower pH. APOL1 inserts into membranes at low pH and demonstrates cation channel activity in *in vitro* and *in vivo* models (10,13,29,44–48), but at least in the presence of DPC we observed no evidence from NMR linewidth or peak intensities, that the C-terminal protein domain of APOL1-G0 forms dimers or higher-order associated states at the lower pH. It is likely that additional protein domains and an extended bilayer membrane model is necessary for this behavior of the protein. The protonation status of D348 within the predicted transmembrane (TM) domain of APOL1-G0 C-terminus (D337-E335) is involved in pH gating function (46) and the G1 and G2 variants demonstrated increased cation channel activity in phospholipid vesicles (48). This predicted TM region in the C-terminus forms an α-helix with similar residue orientation in our NMR model. The changes in HSQC NMR spectra at low pH in our study support the conformational change associated with protonation of D348 in helix-1 and subsequent membrane insertion. Although the membrane topology of APOL1 is not experimentally proven, this conformation could place the helix-2 of C-terminus in the cytosolic domain during intracellular translocation (49). The C-terminal domain of APOL1 is also known to be critical for other biological functions of APOL1, SNARE protein interaction (e.g. with VAMP8, which we demonstrated in our previous study), and *in vitro* vesicle fusion (6,10,13,19,27,28,44,50). Deletion of C-terminal residues V341-L398, which constitutes the α-helical hairpin results in the loss of APOL1 channel activity and these residues by itself can mediate homotypic vesicle fusion (27,44). Consistent with observations from published studies that suggested interaction of APOL1 with membrane and lipid compartments, (6,10,13–15,17,18,44) our studies suggest that APOL1 C-terminus can interact with a membrane environment. Additionally, the amphipathic helical membrane addressing domain (MAD) located N-terminal to the SRA-interacting C-terminal region of APOL1 proteins has been postulated to be a pH sensor and acts as a “hinge” structure between the C-terminus of APOL1 and the rest of the protein (41,51). APOL1 is known to localize to the endosomal and lysosomal compartment (6,18,19,50,52) and exposure to low pH could stabilize the membrane localization (46,47), given the presence of bis(monoacylglycero)phosphate and phosphatidylinositols in the slightly negatively charged endolysosomal membranes (53,54). This conformation can expose the C-terminal region of APOL1 to the cytoplasm, where interaction with SNAREs and other effector proteins is feasible. Existing data suggest that APOL1 interacts with multiple cellular organelle membranes including ER-Golgi network (20,55), mitochondria (14,16,21,56), endo-lysosomal system (6,18,19,50,52) and plasma membranes (13,17,30). Membrane composition and hence protein structure and orientation can differ among these different subcellular environments (57,58).

The models we have presented need to be interpreted with caution because apart from tertiary chemical shifts (such as those arising from the NMR active main chain nuclei of the phenylalanine aromatic rings and charged moieties, such as two histidine residues), no other experimentally derived tertiary restraints enter into the calculations. The higher energies of G1 structures and their variations across the top ten best structure ensemble is likely partly due to a lack of assignments/chemical shift data, particularly around L352, K357 and E379. G2, which has near complete assignments has a much more uniformly predicted helix-2 (αH2). Whether or not this leads to a slight sliding and alternative packing of hydrophobic sidechains at the αH1–αH2 interface would need confirmation from additional NMR experiments, looking at sidechains. However, since our sidechain assignments are incomplete, this is beyond the scope of the current project. Whether the NMR derived structures represent the actual structure with alternate helix-helix packing or at a minimum indicate subtle conformational and dynamics changes or arises because of missing data is not clear at present. The fact that with few exceptions which were noted in the results section above, the NMR chemical shift as well as the relaxation data are highly similar suggests that the structure of the G0, G1 and G2 C-terminal regions is also similar and that the difference may be more subtle, such as the increased stability of the G1 and G2 variants to thermal and chemical denaturation, noted in our previous study (6).

Lastly, we should mention that recent evidence has shown that genetic variations in *APOL1* gene other than the G1 and G2 variants which are inherited together (called *haplotypes*) may contribute to the differential intracellular behavior of APOL1-G1 and -G2 variants compared to G0 (59). Though these *haplotype* associated residues are located far from the variant location in the primary sequence, protein folding, or domain-domain/protein region interactions of full length APOL1 protein, can place these residues spatially close to the C-terminal region of the APOL1 protein where the G1 and G2 variants are located, impacting the protein structure and function. Our study concentrated on the protein region where the variants primarily associated with kidney disease are located and did not take the whole protein into account. In this setting, future studies aimed at understanding the structure and effect of the kidney disease-associated variants in the context of the full-length protein will be critical. In spite of these limitations, our study provides clues to the structure and conformational dynamics of reference APOL1 C-terminal region and suggested strong similarities but also subtle differences of the preferred conformational states and the dynamics for the reference and variant APOL1s in DPC micelles based on NMR chemical shift observations. Our study of the APOL1 C-terminal region structure complements and advances our understanding of APOL1 structure and function (6,26,27,30), further appreciating the results from computational modeling, by reference to experimental information. We also further advance the hypothesis that subtle structural, if not protein dynamics/stability changes induced by APOL1 kidney disease-associated variants could drive intracellular mechanisms in podocytes that predispose to the pathogenesis of kidney disease (22). The C-terminal domain of reference and variant APOL1 could undergo differential structural changes in response to a “second stress” stimuli attenuating the ability of the G1 and G2 variants to initiate and maintain the protein networks necessary to maintain normal podocyte homeostasis. Strategies to modify the differential structural behavior of G1 and G2 variants will guide the development of therapeutic strategies that could halt the progression of APOL1-variant associated CKD.

## Acknowledgments

This work is in part supported by a NIH R01 grant from the National Eye Institute R01EY029169 and previous grants from NIGMS (R01GM073071 and R01GM092851) to the Buck lab.

## Author contribution

S.M. and M.B. devised the project, S.M. with help and advice from A.H. and S.C. carried out the experiments. S.M. and M.B. analyzed the data. S.M., M.B. and J.S. wrote the manuscript.

---

**Supplementary Figure 1:**
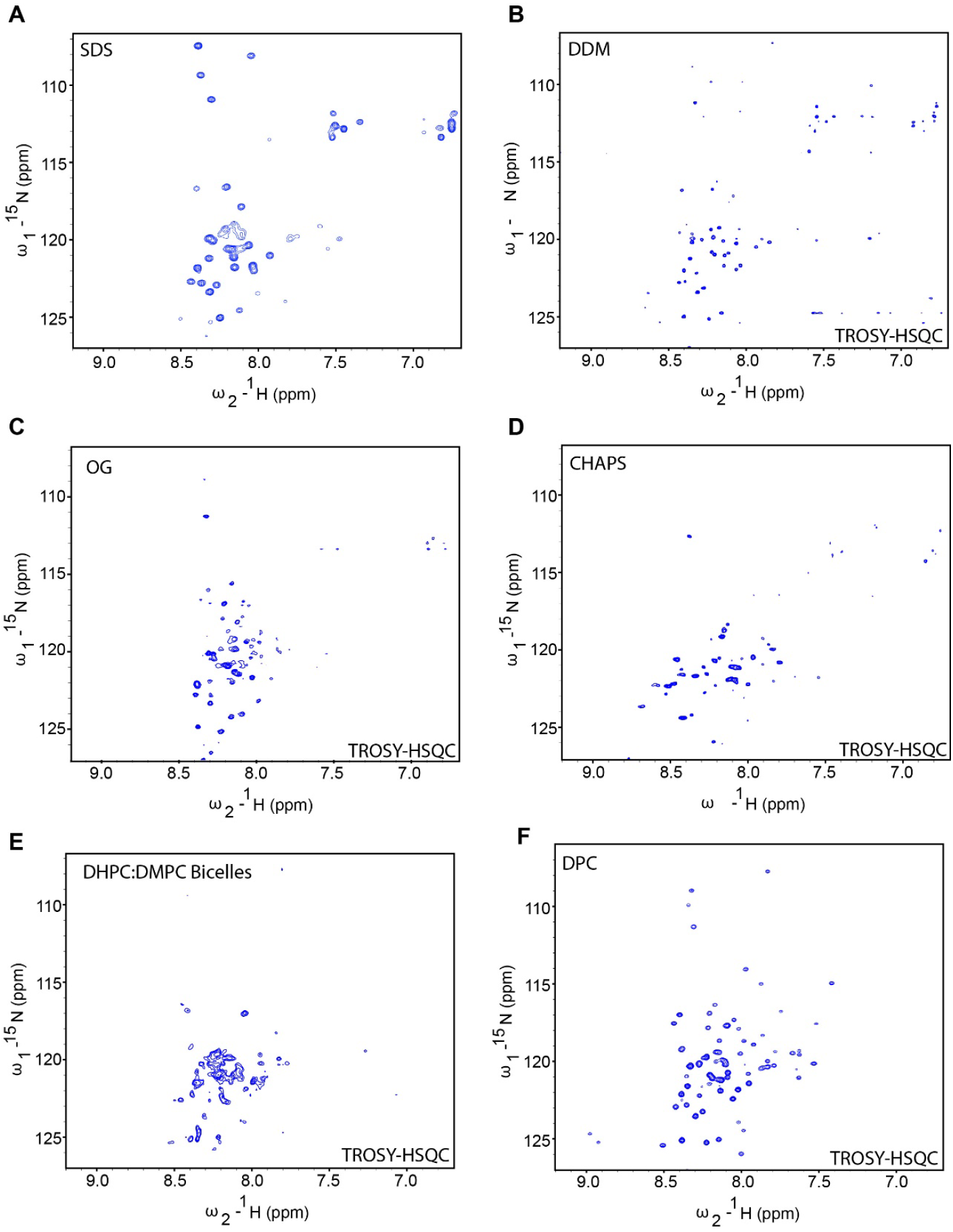
NMR spectra of APOL1-G1 C-terminal in different detergents. ^1^H -^15^N HSQC of domain in (**A**) sodium dodecyl sulfate (SDS), (**B**) n-dodecyl β-D-maltoside (DPC), (**C**) n-octylglucoside (OG), **(D**) 3-((3-cholamidopropyl) dimethylammonio)-1-propanesulfonate (CHAPS) micelles and (**E**) DHPC:DMPC bicelles. Protein concentration was maintained between 0.5-0.6 mM with the concentrations of detergents and bicelles discussed in Methods. The HSQC experiments were TROSY based for all conditions except for in SDS micelles.

**Supplementary Figure 2:**
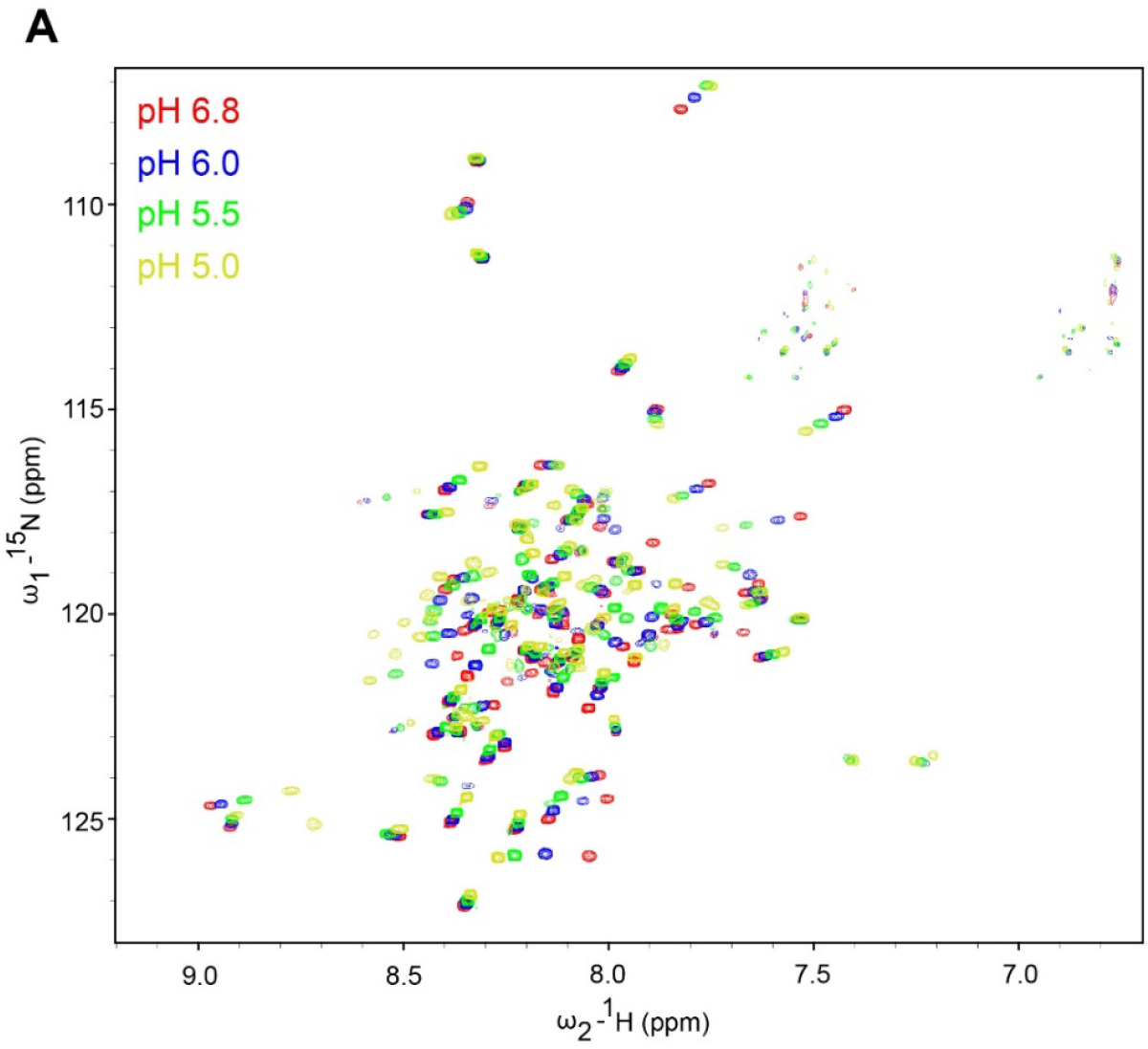
^1^H-^15^N TROSY-HSQC spectra of APOL1-G0 C-terminal domain at different pH conditions. 2D-HSQC spectra in response to varying pH.

**Supplementary Figure 3:**
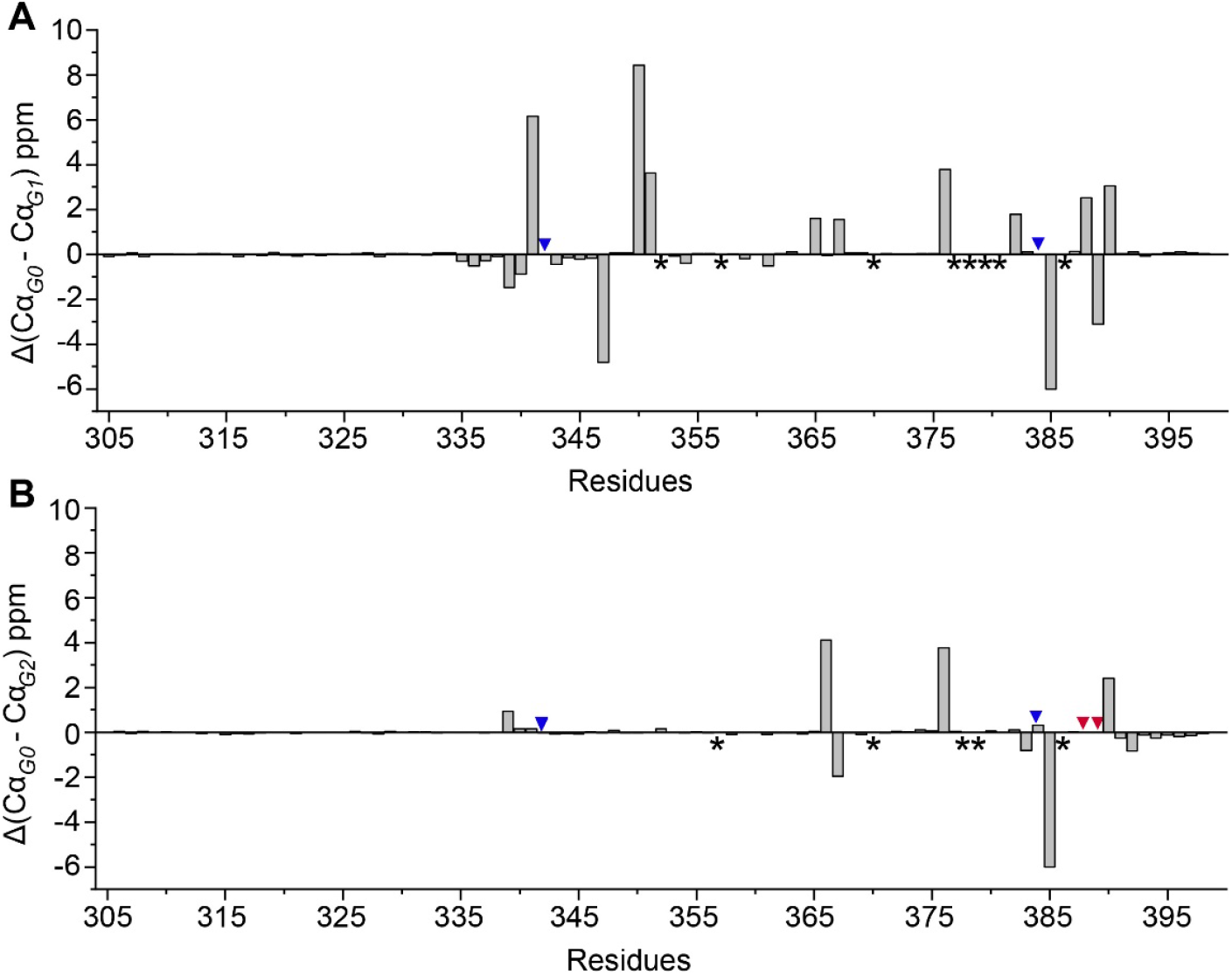
Differences in the Cα chemical shifts of the C-terminal residues of APOL1 G0 and G1 (A) and G0 and G2 (B). Asterisks represents unassigned residues in either G0, G1 or G2. Blue and red arrowheads show the position of G1 and G2 residues respectively.

**Supplementary Figure 4:**
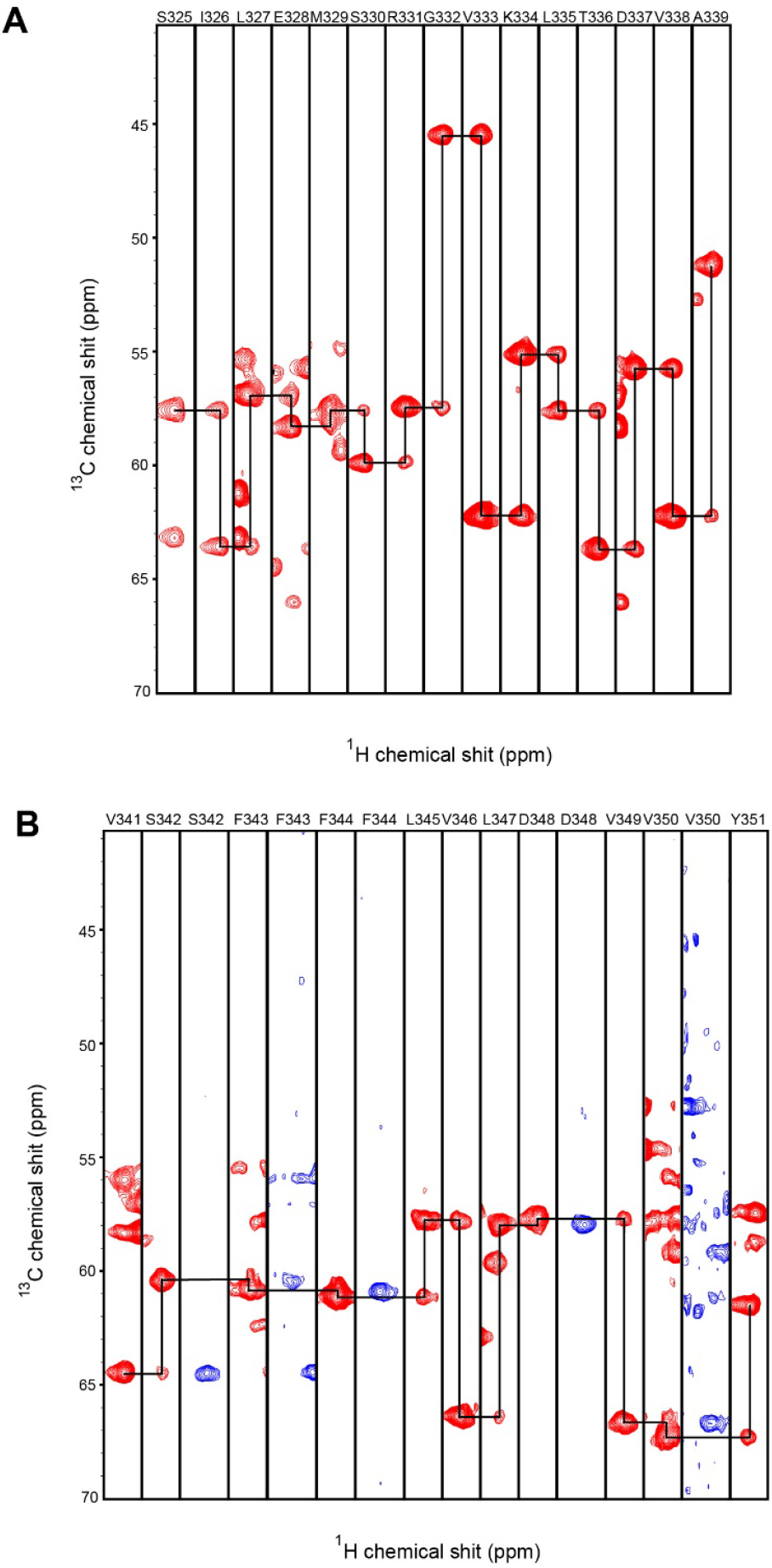
Strip plots of ^13^C chemical shifts of APOL1 G0 from HNCA and HN(CO)CA spectra. Representative plot of two sequential backbone Cα connectivities of (**A**) the N-terminal disordered region and (**B**) the αH1 helix of APOL1 G0 C-terminal region obtained using triple resonance NMR experiments. Lines drawn between adjacent Cα atoms represent the sequential ^13^Cα connectivities for all the residues in G0. The corresponding residues are indicated above the strip plots. The Cα atoms from HNCA and HN(CO)CA are represented in red and blue colors, respectively.

